# Microscopic PhotoSelection (MiPS) of single cells in mother machine microfluidic devices

**DOI:** 10.64898/2025.12.03.692065

**Authors:** Jessica James, Idris Kempf, Kirill Sechkar, Jingyu Wang, Gabriel Abrahams, Sebastian Towers, Harrison Steel

**Affiliations:** Department of Engineering Science, University of Oxford, United Kingdom

**Author notes:** **Equal contribution:** These authors contributed equally to this work (Jessica James, Idris Kempf). **Competing interests:** The authors declare no competing interests. **Data Availability Statement:** The control software used in this work is available on GitHub at https://github.com/kmpape/evomachine/releases/tag/v0.1.0. The modified real-time DeLTA pipeline (DeLTA-RT) is available on GitLab at https://gitlab.com/kmpape/de-lta-rt. The fluorescence image datasets, segmentation masks, and associated retrained models used for ROI identification and cell segmentation are available on Zenodo at doi:10.5281/zenodo.22237153 and doi:10.5281/zenodo.17569655, respectively.

## Abstract

Techniques for selecting or sorting single cells within large populations of genetic variants are central to synthetic biology and biotechnology. Widely-used methods such as Fluorescence Activated Cell Sorting (FACS) enable rapid processing of large libraries, but are restricted to low-dimensional measurements taken at a single time point, preventing their application to dynamic or transient phenotypes. Here we introduce Microscopic PhotoSelection (MiPS) which employs an automated robotic platform for single-cell selection based on multiple dynamic criteria, directly on microfluidic mother machine devices. The system couples long-term single-cell imaging with real-time analysis and selective optical targeting, allowing fully automated screening and enrichment of traits-of-interest. We demonstrate and characterise this capability, and show how iterative, feedback-based selection strategies could refine enrichment across multiple rounds. Taken together, our work establishes a high-throughput methodology to unlock applications of directed evolution and protein design and screening where selection based on dynamic, multi-trait phenotypes is essential.

## Introduction

High-throughput measurement and selection from libraries of biodesigns, in the form of “screening”, underpins contemporary biological engineering. Screening capabilities have been dramatically enhanced by single-cell selection technologies, enabling a shift from traditional arrayed screening— where individual variants are isolated in multiwell plates—to pooled screening approaches, in which millions of variants can be analysed in a single tube. This shift has transformed multiple areas of biology [1]. For example, the vast increases in throughput have been applied for selection-based directed evolution [2] (contrasting growth-coupled directed evolution [3]), which relies on iterative rounds of selection and mutagenesis to alter the properties of a protein.

One of the most widely used high-throughput selection methods is Fluorescence-Activated Cell Sorting (FACS). In FACS, single cells pass at high speed – up to 100,000 per second [4] – through a laser that excites fluorescent molecules and measures a per-cell intensity. Cells are subsequently sorted into separate containers depending on their fluorescence level. An alternative method, Magnetic-Activated Cell Sorting (MACS), allows interrogation of *extra-cellular* markers [5]. In MACS, magnetic beads coated with a target-specific binder (e.g. antibody, lectin) are mixed with the sample, where they bind only to cells expressing the target on their surface. Selection is subsequently performed by passing the sample through a magnetised column. Both FACS and MACS are capable of screening individual cells at high throughput, however they are restricted to single time-point phenotypes (as they only measure each cell once), hence selecting based on multiple and/or dynamic traits remains a major challenge [6, 7]. Where FACS is often applied for intracellular phenotypes, and MACS for surface-based phenotypes, *secreted* phenotypes can be selected upon using droplet-based microfluidics [8]. In droplet-based microfluidics, cells are encapsulated in small droplets that act as microreactors. These droplets can also be combined with various substrates, making this system a suitable tool for the directed evolution of secreted enzymes [9]. Examples exist of droplet based-microfluidics being used to measure dynamic responses by immobilising drops [10], albeit at significantly lower throughput. Long-term observation of cells in droplets presents a challenge due to the finite amount of contained nutrients, as well as the complexity of tracking of single cells in three dimensions.

An alternate family of approaches for high-dimensional single-cell measurements, including during continuous growth/culture, utilises “mother machine” microfluidic devices [11] (Supplementary Fig. S1). The mother machine design consists of a central media channel lined with up to ~ 10^6^ trenches [12], each the width of a single cell. Once loaded into the trenches, the “mother cells” at the closed ends remain effectively static over hundreds of generations, allowing long-term observations. Due to physical separation of cells there is no growth-based competition or selection within the device. In this configuration, dynamic phenotypes can be tracked across many generations at acquisition rates limited only by microscopy and camera measurements (e.g. *>* 100 Hz on standard hardware). Sequential responses to light, chemical, or other environmental stimuli can also be measured and assayed under highly controlled conditions, in ways not possible with other platforms - for example, past work has applied similar techniques for closed-loop optogenetic control of thousands of individual cells over extended periods [13, 14]. To process the resulting data, several computational pipelines have been developed for automated segmentation and tracking of mother machine experiments, making high-dimensional datasets readily accessible [15].

Given the rich phenotypic data that can be collected from mother machine studies, retrieving cells-of-interest (or their associated genetic identity) is of significant interest for unlocking downstream applications. It has been shown possible to link phenotypic and genotypic data in the mother machine using fluorescence *in situ* hybridisation (FISH) [16, 17]. These methods are effective for screening, however they require genetic modifications, extensive sequencing, and in the case of FISH, the cells themselves are destroyed in the process. One approach to extracting cells without the need for genetic modification or destruction is with the use of optical tweezers [18]. This method is able to select a small number of cells from the mother machine, however the process requires a long time period per cell and has only been achieved at low-throughput (e.g. extracting on the order of tens of cells), making it incompatible with high-throughput sorting. It was later shown possible to use strategies such as laser-ablation and optogenetically-induced antibiotic resistance to select cells, paving the way for integration with future high-throughput pipelines [19]. For a high-level comparison of these and other optical screening approaches see Supplementary Table S1.

In this work, we introduce Microscopic PhotoSelection (MiPS) as a method for fully automated, high-throughput selection of cells from mother machine devices using targeted UV light, as well as using targeted red light in combination with the photosensitiser methylene blue. These methods both utilise a Digital Micromirror Device (DMD) to project targeted, patterned light onto cells at single-cell resolution. Using high-power LEDs, we demonstrate it is possible to use the DMD to perform selection on populations in mother machines with *>* 10^4^ trenches, with the potential to expand throughput to the limits of microfluidic device sizes. When compared to FACS—the method currently used for similar challenges—our method unlocks the ability to perform selection based on the rich, multi-factorial data available from mother machine experiments, including dynamic responses [20], tracking of sub-cellular localisation [21], or responses to whole arrays of chemical and/or optogenetic stimuli.

## Results

### Development of Automated Photoselection Platform

To enable high-throughput selection based on dynamic, multi-trait phenotypes we developed an integrated, fully automated platform that couples high-throughput microscopy with microfluidics, real-time data analysis, and targeted optical actuation (Fig. 1 and Supplementary Fig. S1). The platform continuously acquires and processes fluorescence images across multiple fields of view (FOVs), quantifies cell-level traits, and applies selection light patterns without manual intervention (Fig. 1A). Under typical operation (10,000 growth trenches per device; 25 FOVs with 400 trenches per FOV; acquisition every 5 min) the platform must process each FOV within 12 s.

**Figure 1.**
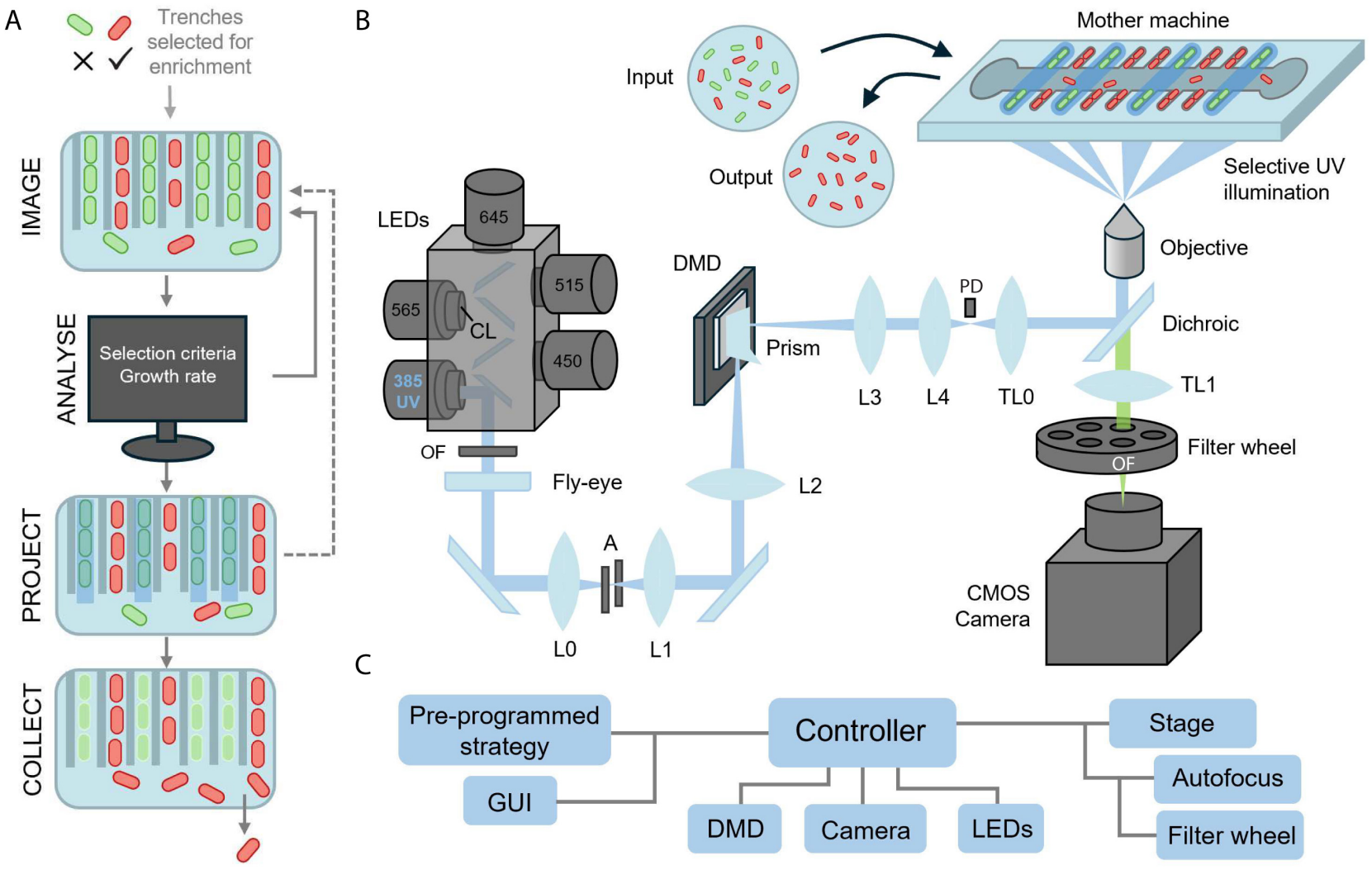
Schematic illustration of the photoselection platform. **A:** Automated pipeline for iterative photoselection. After selecting trenches for enrichment, cells are imaged to compute growth rates, which are then used to generate light projection patterns that selectively kill cells. This process can be iterated to further refine the selected population, after which survivors are collected. **B:** Hardware outline: Light from high-power LEDs (used for both selection and excitation of fluorescent reporters) travels through a fly-eye lens for improved uniformity. A prism positioned directly in front of the digital micromirror device (DMD) ensures that light converges correctly onto the DMD, maximising the area utilised, and ensuring the correct angle of light for reflection into the rest of the optics system. The DMD is an array of micromirrors, which can be individually tilted between two positions (on/off) to dictate the pattern of light projected onto the sample. Patterned light travels via three further lenses and is reflected by the dichroic onto the sample, where it interacts with cells. When imaging the sample, emission wavelength light travels back through the dichroic, stray excitation light is attenuated in the filter wheel, and subsequently detected by the high-speed camera. L = lens, TL = tube lens, CL = collimating lens, OF = Optical Filter, A = aperture, PD = photodiode. **C:** Overview of hardware peripherals controlled by the photoselection platform.

The platform achieves real-time operation by combining automated microscopy hardware (Fig. 1C) with a custom DMD-based actuation system. Throughput is increased by using 40× magnification to maximise the number of cells per FOV and by acquiring fluorescence images only (used for both cell segmentation and phenotype measurement) to minimise data acquisition time. To enable this data to be processed automatically, the DeLTA (Deep Learning for Time-lapse Analysis) pipeline [22] was adapted for real-time operation [23] and the region-of-interest (ROI) identification and cell-segmentation models were retrained on fluorescence images acquired using the 40× objective.

Low-level code optimisation reduced the per-FOV image processing time from 11.6 s to 2.2 s. Training and validation procedures, segmentation performance metrics, and links to the released datasets and models are provided in Supplementary Section *Real-Time Image Processing and Model Retraining*. In routine experiments without applying selection light patterns, the full loop (stage motion plus processing) requires less than 5 s per FOV, below the image-acquisition interval. Another key enabling component is the custom built excitation and actuation light-delivery system (see *Methods*), which allows precise targeting of cells and the duration of selective projections to be reduced. It provides high optical power while minimising off-target illumination, and achieves a uniform light delivery profile across the field-of-view suitable for both quantitative imaging and selective projections.

### Characterisation of Growth Inhibition with Light

To optimise cell selection protocols we began by applying patterned dosing to trenches at different intensities and measuring subsequent reductions in cell growth. In all experiments cell trenches in the mother machine were located and segmented using a modified version of the DeLTA pipeline [22] (Fig. 2A). The coordinates for each identified trench were mapped to DMD coordinates using the calibrated transformation described in Supplementary Section *DMD Calibration* and used to generate the projection pattern. Figure 2B shows a scenario in which UV was projected onto alternating trenches (8.0 W cm^−2^ intensity and 300 s duration), subsequently reducing the division rate of targeted cells to on average ~10 % that of untargeted cells.

**Figure 2.**
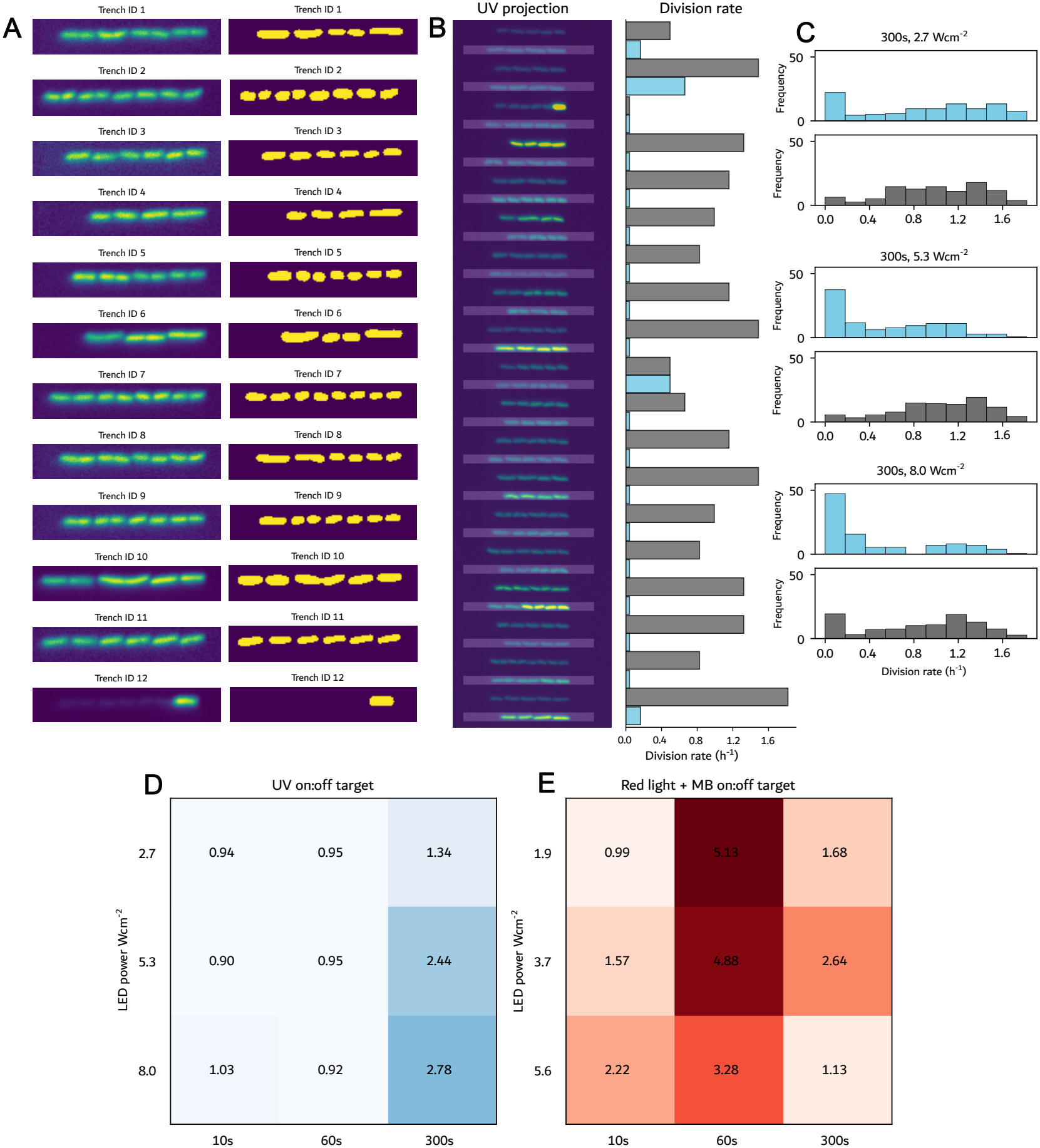
Characterisation of photoselection with UV and red light with the photosensitiser methylene blue (MB). **A**: Segmentation of cells in the mother machine by DeLTA [22], allowing for identification and localisation of cells for UV projection. **B**: Targeting of alternating trenches with UV light projection ≈ 8 W cm^−2^ (90 % of maximum deliverable power) for 300 s (left), and subsequent division rates for each cell (right). Targeted cells’ division rates highlighted in blue, untargeted in grey. **C**: Impact of increasing intensities of UV light on alternating targeted and untargeted cells highlights inter-cell variability in responses. **D**: Relative on:off target killing (higher is better) for UV, and **E**: MB (2 µM) with 645 nm red light.

With the goal of optimising the trade-off between UV dosage and timing, an array of UV intensities and durations was tested (Supplementary Fig. S2). The highest performing selection of conditions tested (300 s duration) is shown in Fig. 2C. Cells with a division rate of less than 0.4 h^−1^ (chosen based on the location of the inter-peak minimum when fitting bimodal distributions to post-UV growth rate data as in Fig. 2C) were deemed to be ‘killed’. After targeting with UV, populations generally exhibited bimodal distributions with respect to division rate (one non-growing population and one with similar-to-initial growth population, and few intermediate rates of growth). This is unsurprising given prior investigations of *E. coli*’s response to oxidative stress [24, 25], which is a key mechanism of cell killing by UV light [26]. Expression of stress response genes in bacterial populations is highly heterogeneous [24], while growth rates stay consistent for a wide range of antioxidant defence enzyme activities before sharply falling for very low levels [25]. As the proportion of targeted cells increases, off-target effects also lead to an increase in the proportion of untargeted cells that are killed; this ratio of on:off target killing is displayed in Fig. 2D. The optimal dose in the tested conditions was 8.0 W cm^−2^ UV for 300 s, which resulted in a 2.78× increase in on:off target ratio with respect to cells killed.

We then explored whether the effectiveness of cell killing could be improved with the use of a photosensitiser chemical. Methylene blue (MB) has previously been shown to cause production of singlet oxygen species in the presence of red light, which can kill *E. coli* in sufficiently high concentrations [27]. In order to test the efficiency of methylene blue, cells in the mother machine were treated with M9 supplemented with 2 µM MB for 1 h, followed by projection with red light (645 nm) following the same protocol as used in the UV dosage experiments (Supplementary Fig. S4). The resultant on:off target killing ratios are displayed in Fig. 2E. By utilising the photosensitiser MB, it was possible to achieve up to 5.13× on:off target killing, a 1.8× improvement over UV, though the fact that the optimum in Fig. 2D lies on the boundary of achievable conditions suggests higher UV dosages may lead to greater killing ratios - though at the expense of a slower selection process. This difference is likely due to the fact that MB has a steeper dose response curve, and thus greater differences between on and off-target effects can be achieved at a given pair of on and off target dosages (Supplementary Fig. S3). In addition, the optimal settings at which this is achieved require lower light doses (60 s and 1.9 W cm^−2^ LED intensity), a 5× reduction in duration relative to the UV optimum. However, it was also observed that addition of MB to the cells causes a growth rate reduction of ≈ 40 % even in cells untreated with red light, and therefore this apparent sensitivity could lead to further issues if subsequent rounds of growth and selection are intended. Meanwhile, while UV is a known mutagen in *E. coli* (via SOS response-mediated induction of error-prone polymerases [28]), MB activates oxidative stress response pathways [29] which (though impacting cell physiology) may avoid induction of SOS [30], supporting its application for situations in which host mutagenesis is undesirable.

### Quantification of Off-Target Effects

From the experiments in Fig. 2 it is clear that off-target effects cause unwanted death of untargeted cells, particularly in the UV setting. Therefore, several experiments were conducted to gain a greater understanding of these effects. First, to understand whether these effects are primarily local (i.e. limited by contrast and accuracy of the projected DMD image or scattering of light from cells to their nearest neighbours) or global (i.e. device-wide scattering or delivery of off-target or un-focused light), an experiment was conducted in which a strip of UV was projected on one or multiple neighbouring trenches, and the division rate of the immediate neighbours assessed (Fig. 3A). It was found that there was no significant difference in division rate between a cell one position away from the targeted trenches versus two positions away (or more), suggesting that off-target effects are *not* primarily caused by local effects arising from blurring of DMD light delivery, and instead occur at a larger or global scale. This provides validation for the design of our DMD and optical system, namely that sufficient local (i.e. pixel to pixel) contrast is achieved in image projection to make this factor insignificant compared to other sources of stray light (e.g. reflection and scattering).

**Figure 3.**
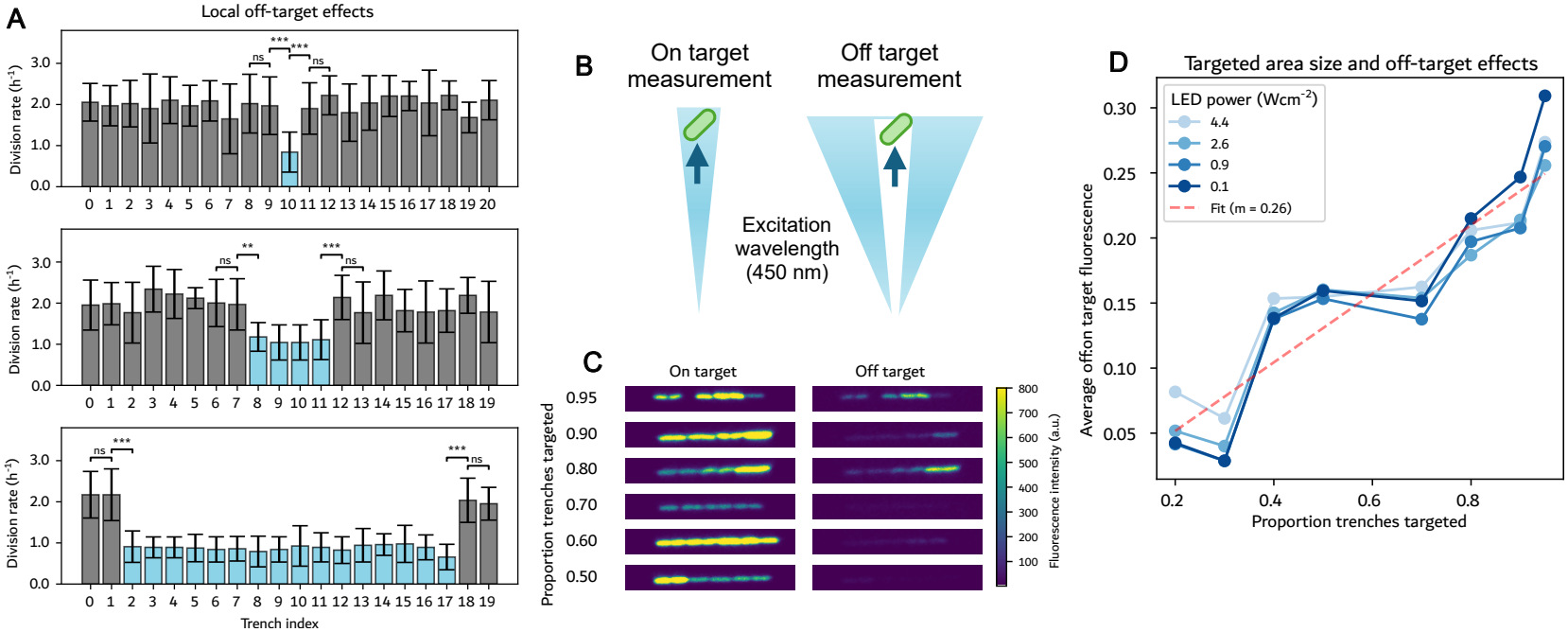
Characterisation of off-target effects. **A**: Experiment to explore local off-target effects, in which an increasingly large strip of UV light was projected down the centre of a sample, and the division rates of the immediate neighbours measured. Targeted trenches are indicated in blue, untargeted in grey. The lack of significant growth slowdown in trenches neighbouring the target region suggests that off-target effects arise from global (i.e. whole field-of-view) rather than local mechanisms. **B**: Schematic of an experiment to explore global off-target effects using the 450 nm excitation wavelength of Green Fluorescent Protein (GFP). An on-target measurement is from illumination of that cell only. An off-target measurement is illumination of an area away from that cell - any fluorescence subsequently collected from that cell is defined as a result of off-target light. **C**: Sample images demonstrating that as the proportion of trenches illuminated increases, the excitation of off-target trenches also increases. **D**: Quantification of the relationship between targeted area size and off-target excitation. Gradient (m) w.r.t proportion cell trenches targeted = 0.26 (90% confidence interval [0.228,0.298]).

A second experiment to quantify the amount of stray light was conducted using selective targeting of fluorescence excitation light (450 nm) to GFP expressing cells, allowing a direct measurement (through fluorescence) of light delivery to each trench (i.e. rather than one mediated by cellular sensitivity to UV measured through growth rate). The assay consists of pairing on-target and off-target measurements of the same trench; in this case an on-target measurement comes from illumination of the target trench (and nothing else), while an off-target measurement comes from illumination of a known area elsewhere in the FOV, and then measuring fluorescence of the target trench arising from excitation by this off-target light (Fig. 3B). The off:on target ratio is then computed by comparing the fluorescence of each trench by off-target light to its on-target fluorescence measurements. A sample of such comparisons is displayed in Fig. 3C. This procedure was conducted across 9 different FOVs, in which an increasing proportion of trenches was illuminated in the off-target tests. 20 individual trenches were measured in each of the FOVs, and the results are shown in Fig. 3D. The ratio of off:on target effects increases approximately linearly with respect to the area of light delivered, and is constant with respect to LED intensity. Off-target excitation was also observed when illuminating regions of the microfluidic chip that do not contain cells or liquid. The exact off:on target ratio expected can be calculated using the proportion of cell trenches targeted, using scale factor 0.26 (the gradient of the linear fit in Fig. 3D). For instance, if 99 % of trenches are targeted, then the untargeted 1 % will receive 0.99 × 0.26 ≈ 25.7 % of the on-target dose. From this data we observe that a large fraction of the light delivered to a given area within the FOV ends up exciting *other* parts of the FOV, likely arising from a combination of scattering in the excitation light path, second order reflections between the cover slip and objective front, and - likely the majority - from reflections and scattering of excitation due to polydimethylsiloxane (PDMS)–glass–media interfaces.

### Implementation of Enrichment by Photoselection

In order to test whether the process of selective UV illumination can effectively lead to downstream enrichment of a desired phenotype, an experiment was set up to test whether mCherry cells could be selectively enriched from a mixed mCherry/GFP population. Immediately before loading into the mother machine, cells were combined in an approximate 1:1 ratio. After loading, cells were allowed to grow for several hours in order to achieve homogeneous trenches (i.e. each trench either contains GFP-*or* mCherry-expressing cells). The mother machine device was then imaged both at 450 nm (for GFP) and 565 nm (for mCherry) excitation, with emission filters complementary to the respective fluorescent proteins. A composite image was used to perform segmentation with DeLTA, and the resulting masks for each trench were compared in both imaging wavelengths to classify trenches as GFP or mCherry (Fig. 4A). Trenches classified as mCherry were selected for survival; all other trenches, including GFP, empty, or unidentified trenches, were targeted with UV by default (Fig. 4B). Details of the fluorescence classification and generation of the DMD projection pattern are provided in Supplementary Section *Real-Time Fluorescence Classification*. UV was projected at 5.3 W cm^−2^ intensity for 5 min (see example in Fig. 4B); the resulting mother cell length traces in Fig. 4C show targeted growth inhibition of GFP cells, indicated by a flat line after the projection at time point 40.

**Figure 4.**
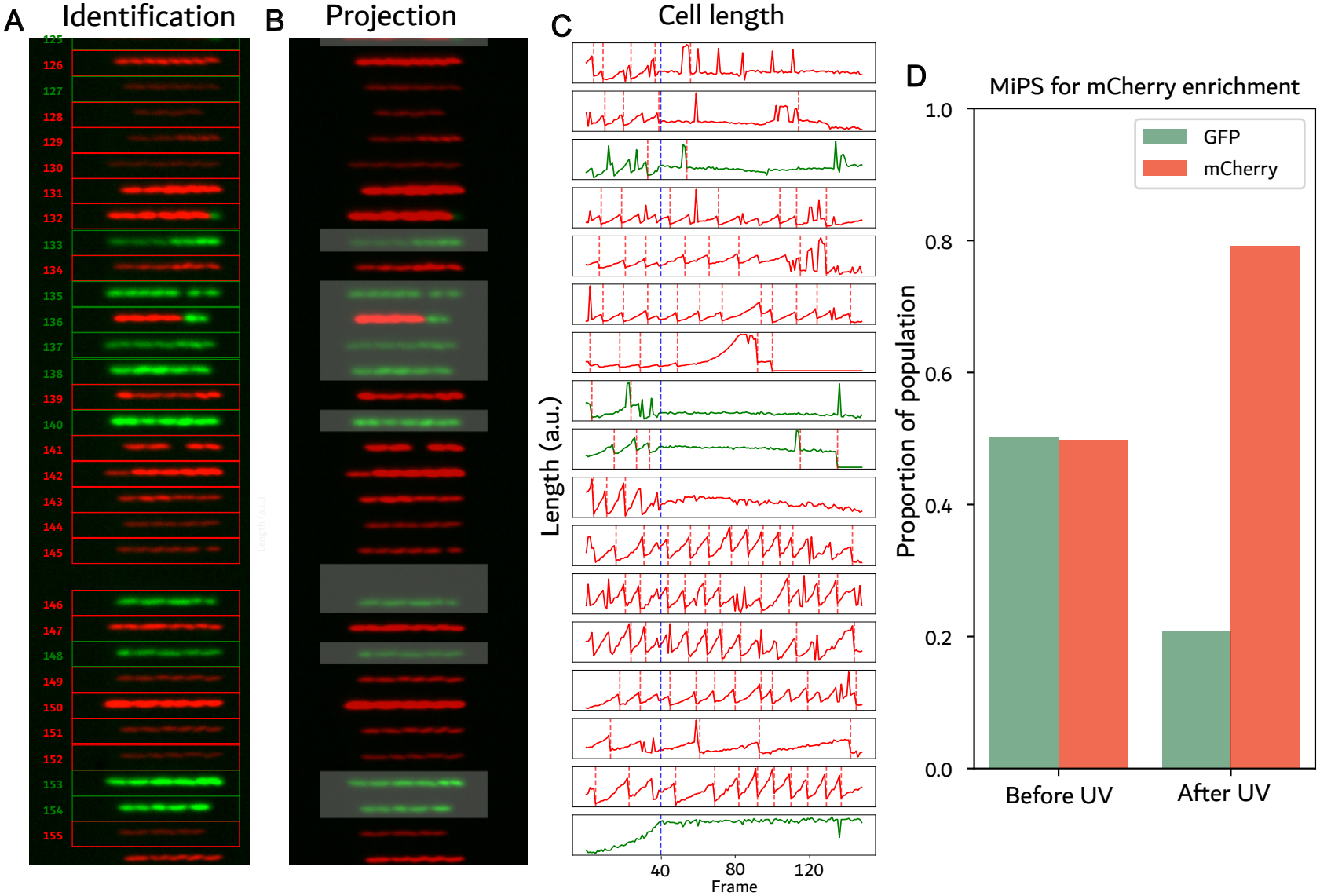
Enrichment of mCherry cells from a mixed mCherry/GFP population by photoselection. **A**:Identification of GFP and mCherry cells was carried out by imaging with a 450 nm LED (GFP, green) and 565 nm LED (mCherry, red), segmenting the images with DeLTA and comparing the brightness in each channel. **B**: The resulting classifications were converted into an image to be projected to target cells for killing. All unidentified or mixed trenches were targeted for killing by default. **C**: Length of the mother cell in each trench can be used to measure divisions (at which point the cell length sharply decreases, as indicated by dashed red line). When UV projection occurs at time point 40 (dashed blue line), the length of GFP-expressing cells (green lines) becomes static, indicating no more division, while mCherry cells (red) continue to divide. **D**: Comparison of the population composition within the mother machine device (pre-enrichment) to the population composition flowing out of the device after UV (post-enrichment; *n* = 1 experiment) shows a reduction in the proportion of the (GFP-expressing) cells targeted for killing.

Following validation that growth could be selectively inhibited on the mother machine device, the population flowing out of the device was analysed. In order to minimise contamination from unselected cells, all microfluidic tubing was replaced with fresh, sterile tubing. The population flowing out of the device was collected for a period of 30 min before being centrifuged to concentrate cells and plated on agar. Figure 4D displays the resulting colony counts from 48 h post-experiment, and shows a shift in population composition from 1:1 mCherry:GFP to 3.8:1 after a single round of targeted selection. In an ideal system *no* targeted cells would survive selection (in which case this ratio would be much larger). However, as illustrated by our characterisation experiments above, achieving this outcome is limited by the off-target delivery – if the UV dose is increased further, it will increasingly kill the desired (non targeted) cells and hence may not necessarily improve the enrichment ratio.

### Simulation of Subsequent Enrichment Rounds

The previous sections illustrate the limitations imposed by off-target effects; below we simulate how these *could* be overcome by implementing multiple rounds of selection to refine the output, i.e. the final ratio of targeted to untargeted cells. Between each round, the division rate of remaining cells is assessed; this can be achieved with 97 % accuracy within ~45 min (Supplementary Fig. S6). For subsequent rounds only cells that were chosen for killing but were *not* killed are re-targeted, thereby reducing undesirable killing of untargeted cells (Fig. 5A). This feedback-based approach succeeds due to fewer cells being targeted in each successive round (e.g. only the survivors), meaning *total* UV dose is reduced each round, preventing off-target effects from killing the desired population.

**Figure 5.**
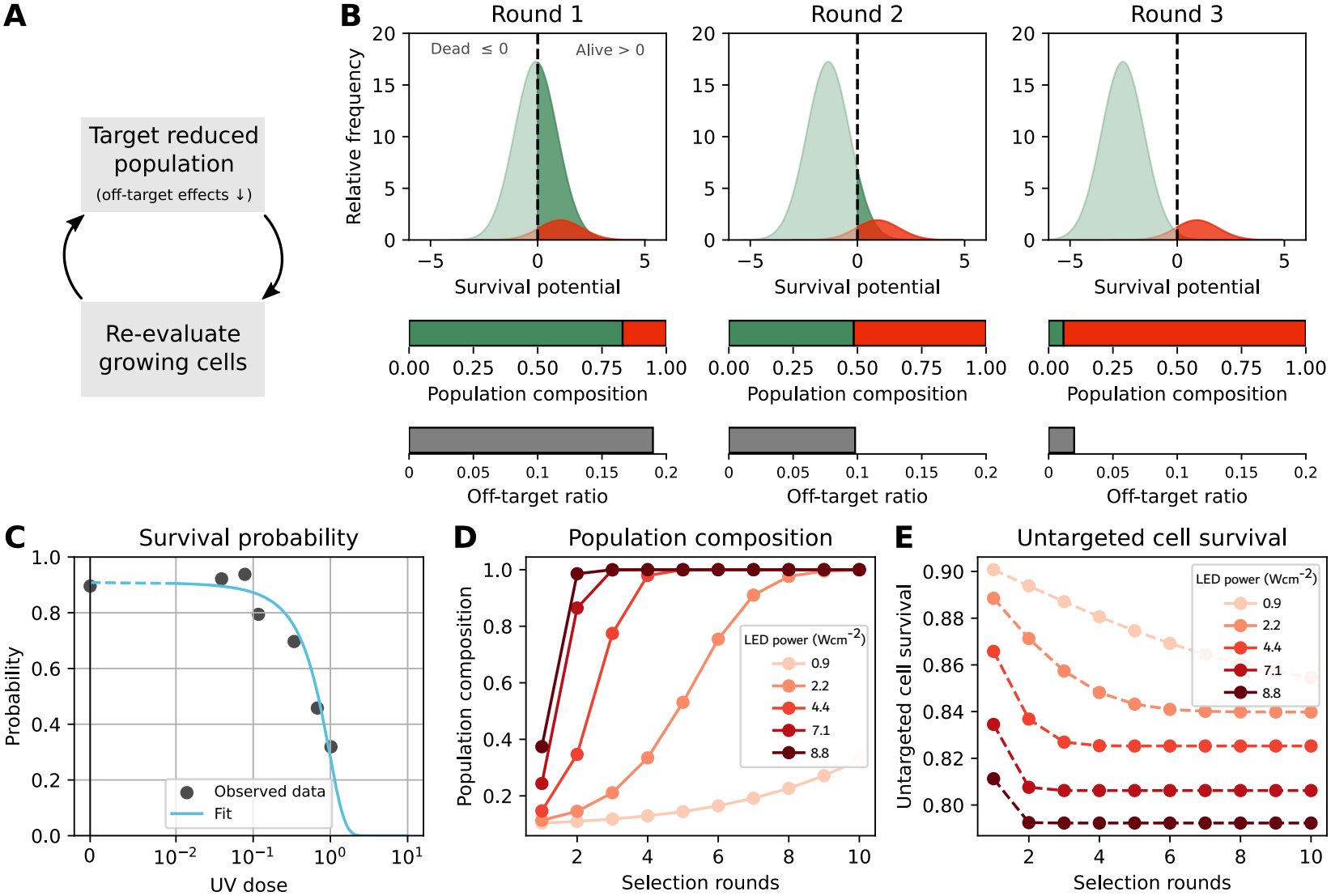
Simulating multiple rounds of targeted selection. **A**: A continuous selection cycle iterates through steps of cell killing and evaluation; off-target effects decrease with each round allowing iterative enrichment of selected population. **B**: Visualising three rounds of simulated selection, including population composition (green = targeted by UV for killing, red = untargeted), and the fraction of light (as a percentage of on-target light intensity) that reaches off-target cells in each round (grey bar). **C**: Survival probability fit from UV dosage experiments (Supplementary Section *UV Selection Model*). One unit of UV dose corresponds to ≈ 8.8 W cm^−2^ for 300 s. **D**: Population composition and **E**: untargeted cell survival over 10 rounds with different UV intensities. Taken together these simulations highlight a trade-off between minimising the number of selection rounds (favouring large LED intensities) and maximising untargeted cell survival (favouring small intensities).

We modelled this iterative process using a probabilistic UV-killing model fitted directly to the dose-response measurements (Supplementary Section *UV Selection Model*) to account for inter-cellular variability in UV sensitivity, which resulted in a sigmoid-shaped survival probability curve as UV dose increases (Fig. 5C). At each round of the simulation (see Supplementary Section *Implementation of Simulation*), targeted cells that are still alive are targeted with constant UV doses, which accumulate across rounds because cells are assumed to be retargeted within 45 min without time for recovery. Off-target UV doses are computed from Fig. 3D. Note that the same approach could be used if cells were allowed to recover, which would result in additional selection rounds to meet the same output refinement.

Figure 5B visualises the output of the simulation for a per-round UV dose of 0.6 (corresponding to 5 min at 5.3 W cm^−2^ UV intensity) and an initial population ratio of 10:1 targeted (green) to untargeted (red) cells (i.e. this simulated experiment aims to enrich red cells, which start in a 10:1 minority). The ratio of alive to dead cells is visualised using the normally-distributed survival potential, where the area over the positive domain corresponds to the fraction of alive cells (Supplementary Section *UV Selection Model*). Given the high proportion of targeted cells, the ratio of off-target to on-target light at the beginning is high, at >10 % (calculated from Fig. 3D). With each round, as the proportion of targeted cells remaining decreases, the amount of off-target light received by untargeted cells becomes smaller, but the targeted population continues to receive *approximately* the same, high dose of UV. This is why the populations appear to move apart with subsequent rounds. After three selection rounds, the fraction of targeted cells remaining in the population is 5.8 % (a total enrichment of ~ 160-fold compared to the initial 10:1 ratio), with 0.5 % and 81.9 % of targeted and untargeted cells still being alive respectively. The fraction of remaining untargeted cells could be increased by reducing the per-round UV dose, at the expense of running additional selection rounds to meet the same output refinement.

This trade-off is further explored in Figs. 5D–E for a starting population of 10:1. With 8.8 W cm^−2^ per-round UV intensity (the maximum deliverable by our system), one could enrich to almost 100 % untargeted cells after two rounds (Fig. 5D), however over 20 % untargeted cells are lost in the process (Fig. 5E). In contrast, with 2.2 W cm^−2^ per-round UV intensity, it takes 8 rounds to reach the same population composition, but results in only 14 % loss of untargeted cells. Therefore, one should consider whether to prioritise minimising loss of cells desired for selection, or experiment time (45 min between rounds for reliably evaluating growth, up to 2 h per round for UV projection).

## Discussion

This work develops Microscopic PhotoSelection (MiPS) - a method for applying selection to microfluidic chips that leverages real-time data processing and actuation afforded by high capability robotics and real-time image processing algorithms. This technique enables selection based on dynamic phenotypes in a mother machine context—features fundamentally inaccessible to state-of-the-art workflows such as FACS or MACS. It is also particularly suited for applications requiring *negative* selection (i.e. elimination of undesired phenotypes) such as screening out non-producing variants of a biomanufacturing process (whose increased growth can confound selection-based assays), whereas *positive* selection methods (also implementable with a similar platform, for example through optogenetic induction of antibiotic resistance [14]) may be more appropriate if aiming to select *one* variant out of thousands.

The current implementation is applied to mother machine microfluidic chips with approximately 10 × 10^3^ trenches across the whole device (~ 400 trenches per FOV). This is a limitation of microfluidic device capacity, which can feasibly be increased to the order of 10^6^ by using multiple devices in parallel or more densely-packed serpentine designs as opposed to the standard linear channel [12]. Serpentine chips have been designed to contain twice as many cells per FOV as the linear designs, and with a lower-magnification objective one could fit up to 1500 cells in a single FOV. While the proof-of-concept selection experiment demonstrated here used 300 s of UV light for selection, it would be possible to decrease this time using higher UV intensities (or using the photosensitiser MB alongside targeted red light) to improve selection throughput.

The observation of bimodal population distributions in our photoselection step is mirrored by prior studies of oxidative stress [24], caused both by UV and red light in the presence of photosensitisers [26, 27]. This is explained by populational heterogeneity immediately *prior* to the onset of adverse conditions, with a fraction of cells in favourable environments expressing more stress-response protein at the cost of slower growth [24]. In the future, our platform’s ability to observe cells over time and measure their growth rates could help identify targeted cells which are stress-resistant. This may allow us to overcome the photo selection trade-offs, for example by delivering a trench-specific, non-uniform light intensity from the DMD in proportion to each cell’s (estimated) stress-resistance, which would minimise total light delivered (and hence off-target effects) while optimising targeted killing. This approach would also minimise the influence of inter-cellular variability in UV resistance that might otherwise confound the desired selection target (i.e. by instead enriching cells based on UV survival probability). Such feedback-based dose shaping can be achieved by streaming grey-scale projection patterns rather than the binary patterns we employed.

While this proof-of-concept work focused on selection of cells based on a categorical variable (mCherry or GFP expression), future development could apply the technique for selection of cells based on a continuous variable (e.g. fluorescence of a variant library). In this, we believe a further advantage over FACS will be unlocked, which is the time over which cells can be observed. Gene expression fluctuates naturally over time (e.g. as a function of cell cycle phase, intracellular resource fluctuations, stochastic noise in gene expression), and thus single time-point measurements are inherently noisy. Observing cells over longer durations allows measurements to be averaged or filtered, giving more accurate phenotyping. Beyond simple gene expression measurements, this system also enables exploration of dynamic cellular responses to environmental and chemical perturbations. Particularly useful in the context of directed evolution of biosensors is the ability to challenge cells with an array of stimuli before performing selection. These stimuli could consist of chemical or media variants flowed in via microfluidics, as well as optogenetic stimuli delivered via the DMD. When applying MiPS to such applications, particularly when the fitness landscapes on which selection is being performed are highly rugged, running the selection process in a regime where some de-selected cells survive each selection round may even *improve* the overall performance of the directed evolution process. This is explored in theoretical studies that highlight benefits of including non-selected cells in subsequent rounds of directed evolution to maintain intra-population variability [31].

Overall, our results demonstrate that high-throughput selection from mother machine microfluidic chips can be achieved by integrating a DMD with automated data processing and control algorithms. While mother machines have proved immensely effective tools for understanding dynamic and long-term cell behaviours at the single-cell level, our approach could also be applied to cells on a standard microscope slide when desired phenotypes are compatible with such a configuration. Taken together, these advantages highlight MiPS as an enabler of high-throughput data collection and multi-parametric directed evolution of diverse properties for engineered biotechnologies.

## Methods

### Microscopy Hardware

The custom built microscopy platform (Supplementary Fig. S1E–F) is based on the Rapid Automated Modular Microscope System (RAMM) from Applied Scientific Instrumentation (ASI). The configuration includes a Nikon PLAN APO *λ*D 40x/0.95 N.A. objective (part no. MRD70470), a Teledyne Kinetix sCMOS Camera, a five-band fluorescence filter set (405/445/514/561/640nm Quinta Band Set, Chroma part no. 89904), and Nikon Ti-2 200 mm tube lens (36 mm clear aperture), resulting in a pixel size of 0.1625 µm × 0.1625 µm and a field of view (for the camera’s 3200 × 3200 sensor of 6.5 µm pixels) of 0.52 mm × 0.52 mm. The stage, filter wheel, and hardware autofocus system (ASI CRISP, augmented with periodic software/camera-based autofocus checks) were driven by an ASI TG-1000 Controller. The light source includes a custom-built synchronisation electronics board (based on the Teensy 4.1 ARM Cortex-M7 microcontroller) and a custom-built LED driver system which can provide up to 70 W of power to each of up to eight LED channels with hardware-based feedback control of LED current or optical output power. The system was configured with five LEDs with wavelength centres at 385/450/515/565/645nm (Luminus Devices part numbers CBM-50X-UV-Y31-FA380-22, SBR-70-B-R75-KG301, PT-39-G-L51, CBT-90-CG-L11-G100, and PT-39-DR-L51-BD100 respectively). LED power was calibrated by projecting a known number of DMD pixels (and hence area) into the microscope FOV which was captured by a ThorLabs PM100D power meter (with wavelength sensitivity calibration set to the middle of each LED’s emission band, or 400 nm in the case of the UV LED).

### Control Software

The process implemented by the control software [32] is shown in Fig. 1A. Cells selected for killing are identified before UV projection (in the case of Fig. 4) based on fluorescence classification (Supplementary Section *Real-Time Fluorescence Classification*), though the platform *also* allows cells to be identified using any arbitrary or dynamic combination of measured traits. The process then iterates between imaging and analysis to compute growth rates. Based on these growth rates, an algorithm autonomously decides which selected cells to target with a fixed dose of UV. In our experimental tests the projection step from Fig. 1A was only executed once, but the control system allows the process to be repeatedly executed in closed loop to improve enrichment performance; such approaches can also be adapted for controlling cell phenotypes (e.g. as in [14]).

Programmatically, this is implemented in a callback-based framework, similar to existing frameworks such as CyberSco.Py [33], MicroMator [34], or Cheetah [35], but low-level optimised and parallelised for throughput. The main process manages image acquisition, hardware peripherals, and data processing. A user-defined *strategy* module—running as an independent process—is provided with processed per-cell measurements and returns high-level instructions (e.g. which cells to target), ensuring that strategy evaluation never interferes with acquisition timing. An optional graphical user interface (GUI)—also implemented as a separate process—allows users to configure strategy parameters.

The main process is implemented in Python on a PC equipped with an NVIDIA RTX 3090 Ti GPU (24 GB VRAM) and 256 GB RAM. It interfaces with the modified high-throughput DeLTA pipeline [22, 23] described in Supplementary Section *Real-Time Image Processing and Model Retraining*, and automates and coordinates the microscopy hardware and software components from Fig. 1C. The camera is controlled using pycro-manager [36]. The stage, filter wheel, and the autofocus system are controlled using a Python serial interface with the ASI TG-1000 Controller (building on the open-source repository [37]). A similar interface is used to connect to the synchronisation electronics board (itself running a custom operating system in C++), which controls the LEDs, reads measurements from a photodiode in the light excitation path if used, and connects hardware triggers. The per-FOV processing time of 5 s is met *without* hardware trigger functionalities, e.g. between LEDs and camera, and usage of this could further reduce processing time.

The DMD is operated using the Video Pattern Mode of the DLP LightCrafter software [38] and connected to the control PC via 4K HDMI. On the control PC, this HDMI connection appears as a second monitor onto which the patterns are displayed using a lightweight display window, implemented in C using Simple DirectMedia Layer (SDL3) and interfaced via a local websocket with the main Python control software. A photodiode placed in the excitation light path can be used to detect the exact moment at which the DMD updates the pattern, allowing fast synchronisation of imaging and actuation if necessary. Additionally, the control software implements a standard software calibration routine that projects a grid of dots on the DMD and records their location on the camera image. It then computes a homography map (specifically, using `findHomography` from the `OpenCV` package [39]) that is used to warp patterns projected on the DMD to account for any misalignment or distortion arising in the post-DMD optics. Further details of the calibration procedure and qualitative validation of projection accuracy are provided in Supplementary Section *DMD Calibration*.

The version of the control software used in this work, including the DMD calibration routines, is publicly available in the EvoMachine release [32]. The modified real-time DeLTA pipeline is available as DeLTA-RT [23], and the retrained models and corresponding training and validation datasets are available through Zenodo [40, 41].

### Optics and DMD Integration

The excitation light delivery system balances the need to deliver large optical power to the sample while minimising off-target and stray light. These design requirements are similar to those of modern DMD-based data projectors, where both large light intensities (a bright image) and contrasts (e.g. brightness ratio of on versus off pixels, often quantified in terms of ANSI contrast) are desirable [42, 43]. Good spatial uniformity of illumination intensity (e.g. across the microscope’s field of view) is also desirable to ensure both quantitative imaging (e.g. fluorescence excitation intensity is consistent from image centre to corner) and uniform dosing when light is delivered to kill cells.

The light delivery system of the photoselection platform is illustrated in Fig. 1B. Excitation LEDs are first collimated using aspheric condenser lenses (Thorlabs ACL25416U-A), and then combined with dichroics with cut-off wavelengths at 600 nm, 525 nm, 470 nm, 425 nm respectively. The combined beam passes through the excitation filter of the five-band filter set (Chroma, described above) before reaching a custom-made two-sided microlens array / fly-eye (16 mm×22 mm external dimensions, 9.5 mm thick, 0.6 mm×1.0 mm microlens dimensions, ~ 5.9° by 12° beam divergence) to create a uniform rectangular illumination profile [44–46] (contrasting the Gaussian-like collimated beam intensity) of approximately the same aspect ratio as the DMD. The fly-eye is placed at one focal length (FL) from lens L0 (180 mm FL Thorlabs AC508-180-A) with a broadband dielectric mirror between (Thorlabs BB2-E02, chosen for good reflectivity at 380 nm). An adjustable rectangular aperture (3D printed) is placed at the focal point between L0 and L1 (200 mm FL Thorlabs ACT508-200-A), allowing the beam shape to be cut to ensure the DMD is not over-filled. This is followed by another lens (L2, 100 mm FL Thorlabs AC508-100-A), with the relative FLs of L1/L2 chosen to create a 4-f relay to de-magnify the fly-eye lens image to fill the DMD’s 8.6 mm by 14.7 mm micromirror array. Light arrives onto the DMD through a TIR Prism [47] (adapted from a large industrial data projector) allowing a setup similar to [48]. Use of a prism here allows closer positioning of the optics L2/L3 (as incoming/outgoing beams require 17.5° angular separation following the DLP670 DMD’s mirror tilt angle), which in turn allows shorter focal-length (and smaller diameter) lenses in these positions and hence collection by L3 (75 mm FL Thorlabs AC508-075-A) of a greater amount of the diffracted beam emerging from the DMD (i.e. part of multiple overlapping diffraction orders, see below). In this setup alignment of L2, DMD face, prism, and L3 are critical as the system must collect the reflected light maxima (emerging approximately normal to the DMD on account of the prism orientation) *and* the maxima of the diffraction that arises from the 5.4 µm pitch micromirrors [49] – and in our setup this would ideally be achieved simultaneously for all five wavelength bands. To facilitate this the prism was mounted on a micrometer X-Y translation stage for precise positioning, and the angle and positioning of the incoming beam was adjusted by translating and rotating the *entire* optical setup upstream of the prism (i.e. including L2 through to the LEDs, all of which are mounted on a separate optical breadboard), to align diffraction bands of order 8 at 390 nm, 7 at 450 nm, 6 at 515 nm, 5 at 645 nm, and mid-way between 5th-6th orders at 565 nm (giving this wavelength the worst diffraction/delivery efficiency) at the focal point after L3. The challenge of maximising transmitted power across multiple diffracted wavelengths is addressed in DMD-based projector and lithography systems via a similar approach [49, 50], though in our setup this is made more difficult by the two additional wavelengths. With L3, L4 (AC508-150-A) is chosen to create a 4-f relay system such that, when combined with TL0 (160 mm FL Edmund Optics # 49-382), the DMD’s image is magnified to slightly over-fill the microscope field-of-view visible to the camera’s 20.8 mm square sensor. A monitor photodiode for DMD synchronisation (described above) is placed at the focal point between L4 and TL0.

Future development of the platform could improve uniformity further by using the DMD’s PWM function (e.g. fast switching of each micromirror) to invert the optical transfer function of the optics between it and the sample e.g. to account for reduced transmission in corners of the objective’s field of view. By comparing the results of our measurements of cells in microfluidics against projection of static patterns on static, flat fluorescent targets (where the system typically achieves >100:1 ANSI contrast, and note that ANSI contrast metrics are highly dependent on the optical setup and are hence typically orders of magnitude less than on-off contrast ratios [51]), we believe the measured off target effects in Fig. 3 are largely due to scattering and other optical effects emerging from the *sample* (i.e. rather than the projection system). However, if improved optical contrast is desired in the future this might be achieved by introduction of apertures at various points (e.g. immediately adjacent to L2 and L3) which can cut out “ghost rays” (second order reflections between the optical elements [52, 53]), which in our setup will be particularly significant in the UVA wavelengths where optic anti-reflective coatings are least effective. However, *in general* there are trade-offs between maximising optical power and contrast for such optical designs [54], and even with further optimisation it may remain challenging to improve ANSI contrast to the levels achieved in commercial data projectors due to the large number of optical elements in microscope objectives which are not necessarily optimised for this function (particularly in UVA wavelengths).

### Microfluidic Chip Fabrication

Polydimethylsiloxane (PDMS) mother machines were fabricated by first combining elastomer and curing agent from the SYLGARD 184 Silicone Elastomer Kit (Dow: 4019862) in a 10:1 ratio. The mixture was thoroughly mixed with an electric whisk, and poured onto a silicon wafer. The mixture was degassed in a vacuum until all bubbles were removed, and then cured for 2.5 h at 65 °C. The cured PDMS was removed from the wafer, devices cut with a razor blade, and inlet and outlet holes punched with a 0.75 mm hole punch (Darwin Microfluidics: PT-T983-07-25). The final PDMS mother machine was prepared for bonding by removing dust with Scotch tape (3M: 7100235316), washing twice in isopropanol and drying with air. High-precision coverslips (Thorlabs: CG15KH1) were prepared for bonding by sonicating in acetone for 10 min, washing twice with deionised water, then sonicating in isopropanol for 10 min. To bond mother machines to coverslips, both were treated in a Henniker HPT-100 plasma oven at 50 % power, 0.3 mbar for 30 s. Bonding was performed by transferring each coverslip treated face up to a hot plate at 80 °C, then placing the mother machine face down on top and applying even pressure. Bonded chips were baked at 95 °C for 1 h. The mother machine design used (a gift from Stephan Uphoff) has trenches with a 1.2 µm by 1.2 µm square profile and 25 µm length, and 5 µm lateral separation between centre points of each trench.

### Media and Strains

All strains used in this work were MG1655, transformed with pVS-01-01-mCherry (a constitutive mCherry expressing plasmid) or pVS-01-03-eGFP (a constitutive eGFP expressing plasmid). All M9 media (Formedium: MMS0102) used in this work was supplemented with 50 µg mL^−1^ Carbenicillin (Formedium: CAR0005), 0.4 % glucose (Formedium: GLU03), 2 mM MgSO_4_ (Sigma Aldrich: M7506-500G), 0.1 mM CaCl_2_ (Sigma Aldrich: C5670-500G), 0.2 % casamino acids (Formedium: CAS03), 0.2 g L^−1^ Pluronic F-127 (Sigma Aldrich: P2443-250G), 50 mg L^−1^ EDTA Na_2_ ⋅ 2 H_2_O (Sigma Aldrich: E4884-100G), 30.8 µM FeCl_3_ (Sigma Aldrich: 157740-100G), 6.16 µM ZnCl_2_ (Sigma Aldrich: 208086-100G), 0.76 µM CuCl_2_ ⋅ 2 H_2_O (Sigma Aldrich: 307483-500G), 0.42 µM CoCl_2_ (Sigma Aldrich: 232696-100G), 1.62 µM H_3_BO_3_ (Sigma Aldrich: B0394-500G) and 0.081 µM MnCl_2_ ⋅ 4 H_2_O (Sigma Aldrich: 221279-500G). For Methylene Blue (MB) experiments, the same M9 media was supplemented with MB at 2 µM in the mother machine, which (without light delivery) was observed to cause an approximately 40 % reduction in growth rate. Independently, we tested the impact of different MB concentrations on exponential growth rate of the same strains in a Tecan Spark plate reader. Here it was found that a 40 % reduction in growth occurred at approximately 16 µM concentration, a factor of ~ 8 greater than that required to achieve the same effect in the microfluidic measurement. Such quantitative differences between experimental systems suggest that future protocol optimisation may be best done in the microfluidic application environment [55], rather than standard well-plates in which optimal conditions may differ significantly.

### Microfluidic Experiments

Overnight cell cultures were prepared in 5 mL M9 media. The following day, mother machines were injected with 15 µL of 8.5 mg mL^−1^ Pluronic F-127 (Sigma Aldrich: P2443-250G), using a gel loading tip (Fisher: 11927734). After approximately 10 min, the overnight cell cultures were injected into the chip in the same manner. In order to load cells into trenches, the chip was spun in a centrifuge at 3000 rpm for 3 min on each side, at an angle of 10°. This was achieved using a 3D-printed chip holder designed to be compatible with a 96-well plate centrifuge rotor (Supplementary Fig. S1C-D).

The loaded chip was then mounted on a custom-designed microscope stage insert that supports the cover slide around all edges (Supplementary Fig. S1E). To insert needles for microfluidics, a 3D printed support was designed that provides a backing for the chip to prevent cracking (Supplementary Fig. S1F). Once connected to the microfluidic pressure flow control system, M9 media with 25 % of the standard antibiotic concentration (12.5 µg mL^−1^ Carbenicillin) was flowed continuously at a rate of 20 µL min^−1^. The pressure is controlled by an OB-1 MK3+ Flow Controller (Elveflow) which delivers pressurised air into a 50 mL reservoir containing M9 media. As a result, media is pushed out into the system, which consists of an MFS Microfluidic Thermal Flow Sensor (Elveflow) which provides flow measurements for feedback control, followed by a 5 cm long, 100 µm inner diameter resistance element (Darwin Microfluidics: LVF-KTU-05) for more robust flow control. After the resistance element, a 0.2 µm filter is attached to prevent contamination reaching the chip or output. From the filter, the tubing connects directly into the chip via glass capillary tubes (DWK Life Sciences: 764500-0002). Cells were maintained at 37 °C by a custom, blacked-out microscope incubator (Supplementary Fig. S1G-H), and allowed to grow for several hours before initialising the experiment.

## Acknowledgment

IK and HS were supported by the Engineering and Physical Sciences Research Council (EPSRC) under the Efficient Engineering and Control of Predictable and Reliable Biosystems (EEBio) Programme Grant, EP/Y014073/1. HS was additionally supported by EPSRC grant EP/X017982/1 and BBSRC grant BB/W012642/1.

