## Supplementary Material for "Microscopic PhotoSelection (MiPS) of single cells in mother machine microfluidic devices"

### Real-Time Image Processing and Model Retraining

For image processing, we modified the DeLTA pipeline [1] and adapted it for real-time operation [2]. DeLTA was developed for post-processing and operates in three stages, each using a specialised U-Net. First, a reference frame from each field of view (FOV) is used to identify regions of interest (ROIs), corresponding here to growth trenches. This stage returns one cropping box per ROI, which is used to extract per-ROI time-lapse image series. Second, each per-ROI frame is segmented to obtain cell masks. Finally, the per-ROI images and segmentation masks are used to track individual cells between frames.

The original DeLTA implementation processes each ROI over multiple time points and communicates with the GPU ROI-by-ROI. In contrast, real-time operation requires all ROIs from a single time point to be processed within the image-acquisition interval. With the original implementation, this creates substantial GPU communication overhead. We therefore made two modifications. First, all ROIs within one FOV are grouped and sent to the GPU as a single batch. Second, we replaced the third, U-Net-based tracking stage with a rule-based algorithm that tracks only the mother cell at the closed end of each trench. This is possible because the geometry and orientation of the microfluidic chip are known. All other features of the original DeLTA pipeline, such as drift and rotation correction, were left unmodified. The modified pipeline is available on GitLab [2].

In addition to modifying the pipeline, we retrained the U-Nets for ROI identification and cell segmentation using fluorescence images acquired with a 40× objective. In contrast, the original DeLTA models were developed primarily for phase-contrast or bright-field images acquired with 100× objectives. The default DeLTA mother-machine models use input sizes of $512 \times 512$ pixels for ROI identification and $256 \times 32$ pixels for cell segmentation, five U-Net levels, and $3 \times 3$ convolution kernels. We retained five U-Net levels and the default numbers of convolutional filters per level, but used $5 \times 5$ convolution kernels and input sizes of $200 \times 800$ pixels for ROI identification and $64 \times 512$ pixels for cell segmentation.

The ROI training data combined manually annotated images of cells grown in LB and M9. ROI masks were manually annotated for the first frame of each FOV and applied to the following 12 frames, acquired at 10 min intervals, after confirming that drift remained within acceptable bounds. The mixed dataset contained 14,014 images, of which one lacked a corresponding mask and was excluded. The remaining 14,013 image–mask pairs were divided into 13,312 training and 701 validation images using a 95:5 split. The model was initialised from an LB-trained model and retrained on the mixed dataset for 500 epochs, using 500 steps per epoch and a learning rate of $10^{-4}$.

The cell-segmentation dataset contained 4,344 manually annotated image–mask pairs, divided into 3,475 training and 869 validation images using an 80:20 split. The segmentation model was trained for 400 epochs, using 1,000 steps per epoch. No independent test set was available for either model. The datasets and models are available under [3] and [4].

On the ROI validation set, the retrained model achieved a macro Dice score of $0.79 \pm 0.18$ and a macro IoU of $0.68 \pm 0.15$. Following the standard DeLTA post-processing procedure, 3,541 of 4,073 annotated ROIs were correctly identified (86.9%), while 91.6% of predicted ROIs corresponded to an annotated ROI. On the cell-segmentation validation set, the model achieved a macro Dice score of $0.747 \pm 0.231$ and a macro IoU of $0.636 \pm 0.216$. In total, 3,236 of 4,068 annotated cells were correctly identified (79.5%), while 87.0% of predicted cells corresponded to an annotated cell. For these object-level measurements, annotated and predicted objects were matched one-to-one and considered correctly identified when their mask IoU was at least 0.5. To compensate for likely mis-identification of trenches based on these model performance metrics we designed the selection protocol to target (for killing) any ambiguous trenches by default.

### Real-Time Fluorescence Classification

To implement fluorescence-based enrichment by photoselection, we implemented the following strategy in the control software [5]. During initialisation, the modified DeLTA pipeline [2] was used to identify regions of interest (ROIs), corresponding to individual growth trenches. Since only fluorescence images were acquired and the mother machine devices were loaded with both GFP- and mCherry-expressing cells, a reference image for segmentation was generated by averaging intensity-normalised images acquired using 450 nm and 565 nm excitation. In the control software, this procedure is automated by specifying the fluorescence channels to be combined for segmentation. Following a predefined imaging period of 2 h, after which each trench was assumed to contain either GFP- or mCherry-expressing cells, trenches containing mCherry-expressing cells were selected for survival. We evaluated several classification criteria, including mean channel intensity and percentile-based thresholds, and found that classification based on the maximum intensity within each trench in the non-normalised 450 nm and 565 nm images yielded the most reliable separation between GFP- and mCherry-expressing trenches.

Following fluorescence classification, the strategy generated a binary projection pattern for display on the DMD, with UV illumination applied to all regions except those containing cells selected for survival. To construct this pattern, a $3200 \times 3200$ pixel binary mask matching the dimensions of the camera image was first generated. Pixels corresponding to trenches selected for survival were marked, together with an additional 10% spatial margin around each trench to protect against small segmentation or projection errors. The resulting mask was then inverted such that the protected regions remained unilluminated while all remaining regions were exposed to UV. Finally, the projection pattern was geometrically warped using the DMD calibration map and displayed on the DMD during photoselection. All of these functionalities are implemented in the control software [5].

### DMD Calibration

To spatially align DMD projections with the camera image, a calibration routine implemented in the control software [5] was used to determine the mapping between DMD and camera coordinates. Circular patterns with a radius of 5 DMD pixels were sequentially projected on a regular grid with spacing of 150 DMD pixels and an image acquired. The corresponding camera coordinate was identified from the position of maximum fluorescence intensity, with points below an automatically determined intensity threshold discarded. To determine this threshold, an on-screen reference intensity was first calculated as the mean of the maximum image intensities obtained from nine DMD points arranged on a $3 \times 3$ grid and chosen to lie within the camera FOV. An off-screen reference intensity was obtained by projecting a DMD point that lies outside the camera FOV. The acceptance threshold was set halfway between the off-screen and on-screen reference intensities. The resulting pairs of DMD and camera coordinates were used to compute a homography mapping between the two coordinate systems using the `findHomography` function from OpenCV [6], which was subsequently used to warp all projection patterns before display on the DMD.

Ideally, this calibration can be performed directly on the mother machine device. However, since the experiments in this work relied exclusively on fluorescence imaging, no suitable transmitted-light image of the device was available for calibration. The calibration was therefore performed at the same imaging plane using a slide containing fluorescent dye, but without a mother machine device.

Projection accuracy was additionally validated qualitatively after calibration. First, using the fluorescent slide, a $3200 \times 3200$ pixel test image was generated containing four columns of 90 rectangular regions, with each rectangle chosen to match the dimensions of a growth trench. The image was warped using the calibrated homography, displayed on the DMD, and imaged with the camera. The contours of the corresponding rectangles in camera coordinates were then overlaid on the acquired fluorescence image, and each projected region was visually inspected to verify that the full contour was illuminated. This was the case throughout the majority of the FOV, with the exception of approximately the first 2–3 trenches at the top and bottom of each image, where the projected pattern became slightly blurred. The same validation procedure was subsequently repeated using an actual mother machine device, and the observed projection accuracy was consistent with that obtained using the fluorescent slide.

### UV Selection Model

To enable simulations of an iterative selection process (Section Simulation of Subsequent Enrichment Rounds), the UV killing process including off-target effects (Section Quantification of Off-Target Effects) was modelled using a stochastic framework, reflecting the cell-to-cell variability in UV tolerance. First, division rates obtained from a series of UV dosage experiments (Fig. S5A) were used to produce dose-dependent survival probabilities $P_{alive}(D)$ (Fig. S5B), where in Fig. S5A cells with division rates smaller than 0.4 $h^{-1}$ were considered to be dead. The UV doses were adjusted for off-target effects using the fit from Fig. 3.

To extrapolate to higher UV dosages, a curve was fit to the survival probabilities from Fig. S5B as follows. The sigmoid-shaped curve was obtained from the cumulative distribution function (CDF) of a normally distributed survival potential, $X \sim \mathcal{N}(\mu(D), \sigma)$, where $\mu(D)$ is a dose-dependent mean. If $X < 0$ for a given UV dose, the cell is assumed to be dead (Fig. S5C). For each dose $D$, the mean $\mu(D)$ was computed such that the area to the right and left of the origin correspond to $P_{alive}(D)$ and $P_{dead}(D) = 1 - P_{alive}(D)$, respectively. Using the CDF of a standard normal distribution $\Phi(Z)$, $Z \sim \mathcal{N}(0, 1)$, the means in Fig. S5D were obtained from

$$\mu(D) = -\sigma\Phi^{-1}(1 - P_{alive}(D)). \tag{1}$$

A linear curve was then fit to the survival potential means, relating the UV dose to the survival potential mean as

$$\mu(D) = aD + b, \tag{2}$$

where $a = -1.915$ and $b = 1.331$. The curve from Fig. S5B is then obtained from evaluating

$$P_{alive}(D) = 1 - \Phi(-\frac{\mu(D)}{\sigma}), \tag{3}$$

for $D \in [0, 10]$. Note that the standard deviation $\sigma$ is arbitrary and only determines the relative distance between $\mu(D)$ for each $D$, so that $\sigma = 1$ is used throughout.

### Implementation of Simulation

To simulate subsequent selection rounds, we implemented a simulation based on the UV process model (Supplementary Section *UV Selection Model*) and the survival probabilities from Fig. S5B in Python. At each iteration $i$, accumulated UV doses are updated as

$$D^i_{\text{on}} = D^{i-1}_{\text{on}} + \left(1 + mP_{alive}(D^{i-1}_{\text{on}})F^0_{\text{on}}\right) D, \tag{4a}$$

$$D^i_{\text{off}} = D^{i-1}_{\text{off}} + mP_{alive}(D^{i-1}_{\text{on}})F^0_{\text{on}} D, \tag{4b}$$

where $D^i_{\text{on}}$ and $D^i_{\text{off}}$ are doses accumulated by on-target and off-target populations, $m = 0.26$ models off-target effects (Fig. 3D), $D$ is a constant dose delivered at each round, $F^0_{on}$ is the initial ratio of targeted to untargeted cells, and $P_{alive}(\cdot)$ is computed from the survival probability model (Fig. S5B). After $N$ selection rounds, the composition of alive on-target and off-target cells (Population composition in Fig. 5D) is then obtained from $F^N_{\text{on}} = P_{alive}(D^N_{\text{on}})$ and $F^N_{\text{off}} = P_{alive}(D^N_{\text{off}})$. Figure 5B also shows the off:on-target UV dose ratio computed from $mP_{alive}(D^i_{\text{off}})F^0_{\text{on}} / \left(1 + mP_{alive}(D^i_{\text{on}})F^0_{\text{on}}\right)$. The assumption that UV-induced damage is *cumulative* would depend on the balance between cellular recovery rate and time between successive doses, as well as whether highly-UV-tolerant cells at one time point *remain* tolerant at later time-points (e.g. as would be the case if tolerance was determined by genetic differences, which would be enriched by our selection process) or if the tolerance distribution is essentially “reset” between killing rounds (e.g. if tolerance variability arises from short-term phenotypic variation and noise).

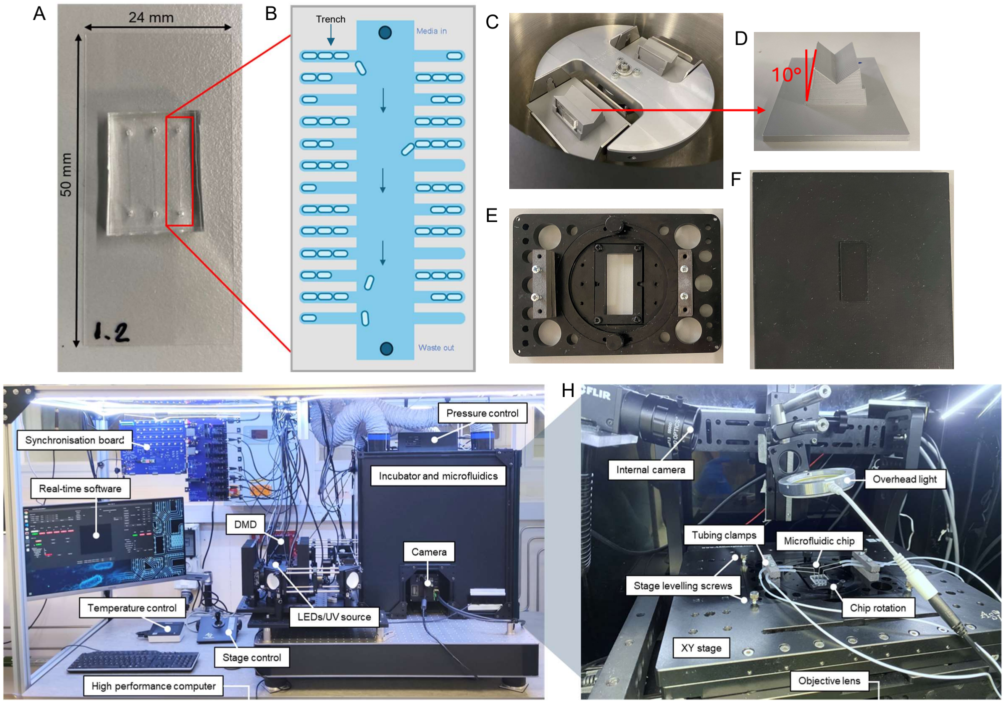


**Figure S1. System hardware**. **A:** Mother machine microfluidic chip. **B:** Mother machine schematic. **C:** 96-well plate centrifuge rotor with 3D-printed chip holder for mother machine loading. **D:** 3D-printed chip holder, with 10° angle. **E:** Custom microscope stage insert with tube clamps, rotational capability, and levelling screws. **F:** Support compatible with stage insert, allowing needles to be inserted into the chip without causing bending (and cracking) of the chip. **G:** External hardware overview including control computer, syncboard, stage control, microfluidic control and temperature-controlled, blacked-out incubator. **H:** Internal hardware including the custom microscope stage insert, with chip connected to microfluidics. An overhead light can be used for bright field imaging.

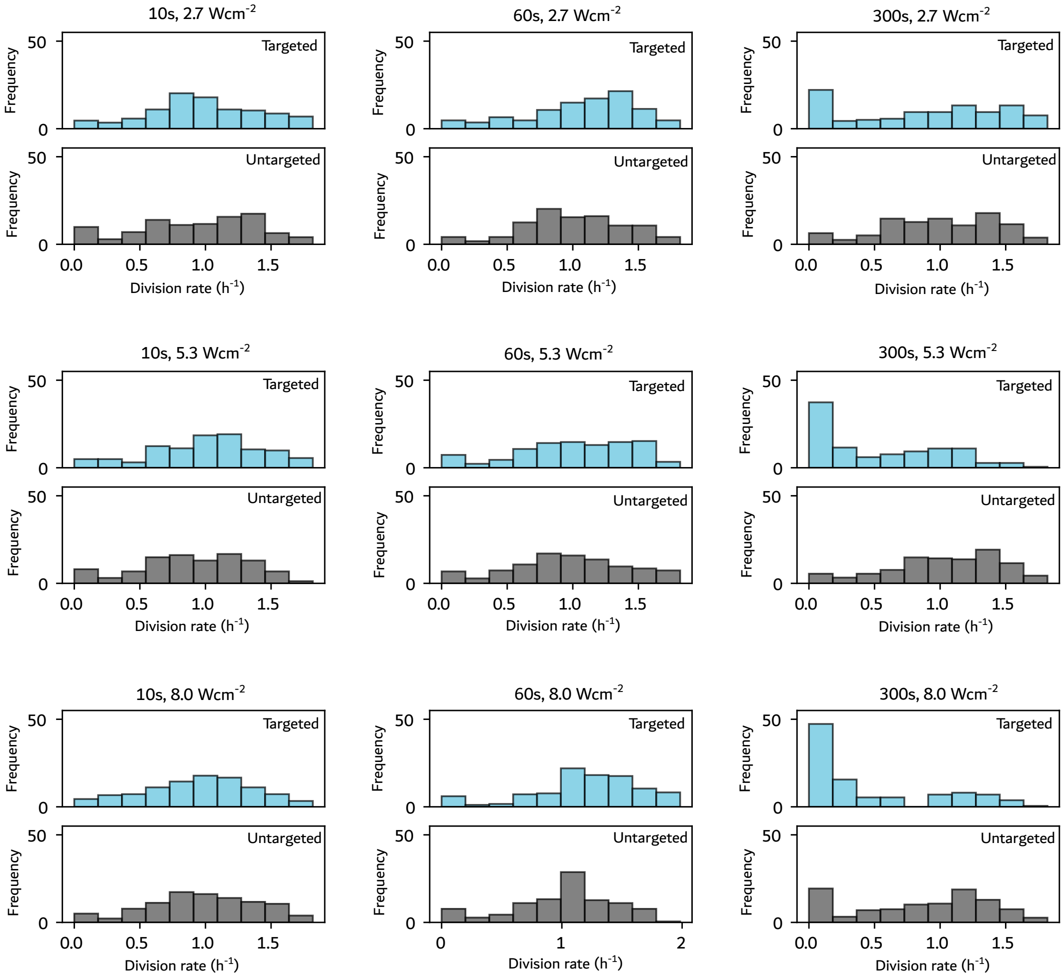


**Figure S2. UV dosage testing**. Alternating trenches in the mother machine were targeted with UV light (385 nm) at the intensities and durations above. The subsequent division rates of targeted (blue) and untargeted (grey) cells are compared.

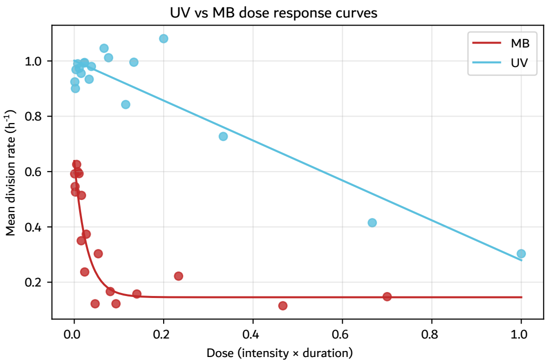


**Figure S3. A comparison of dose response curves between UV and red light with methylene blue (MB)**. For MB, cells were first treated with 2 μM methylene blue for 1 h, then alternating trenches in the mother machine were targeted with red light (645 nm). No prior treatment was required for the UV case, in which alternating trenches were targeted with UV light (385 nm). Dosages have been adjusted for the difference in LED power (90 % 385 nm ≈ 8 W cm$^{-2}$, whereas 90 % 645 nm ≈ 5.6 W cm$^{-2}$).

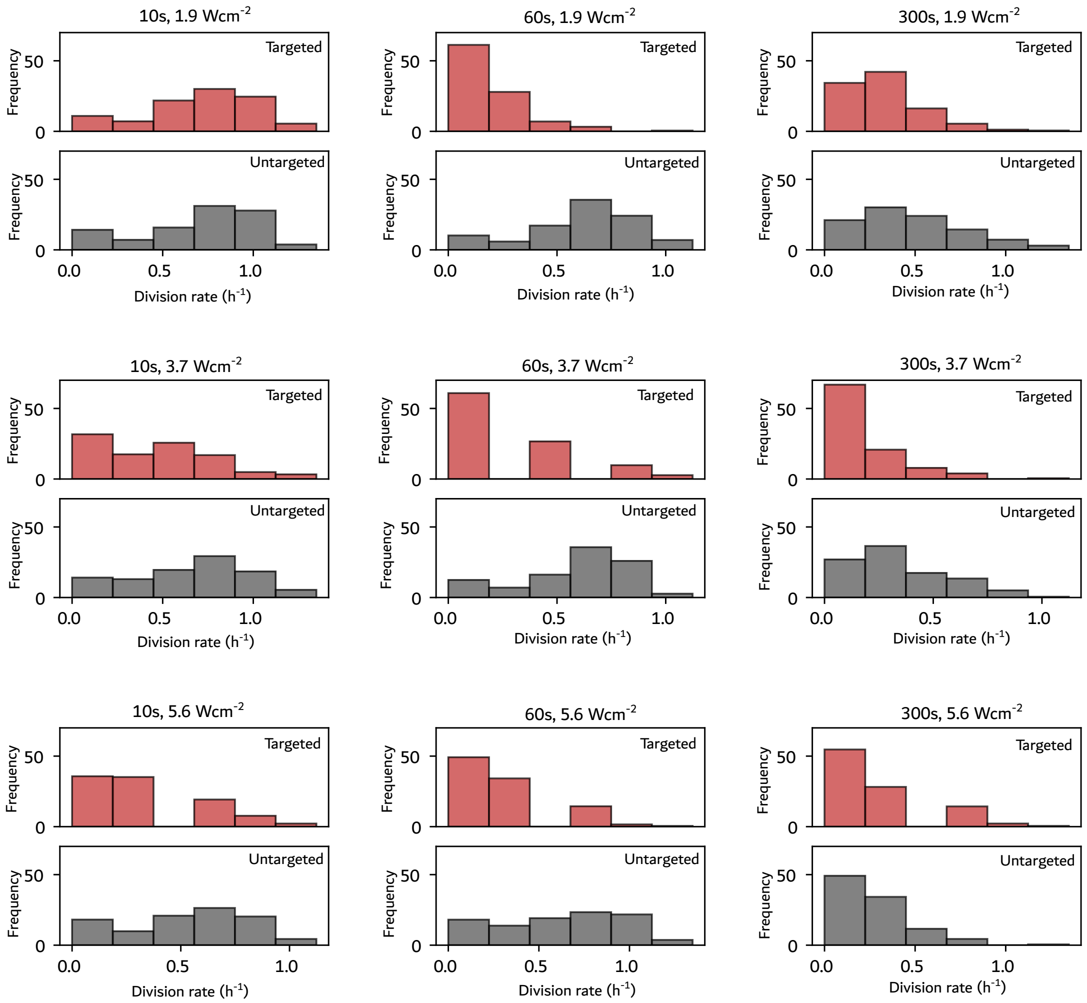


**Figure S4. Red light and methylene blue dosage testing**. Cells were first treated with 2 µM methylene blue for 1 h, then alternating trenches in the mother machine were targeted with red light (645 nm) at the intensities and durations above. The subsequent division rates of targeted (blue) and untargeted (grey) cells are compared. 90 % intensity $\approx$ 5.6 W cm$^{-2}$.

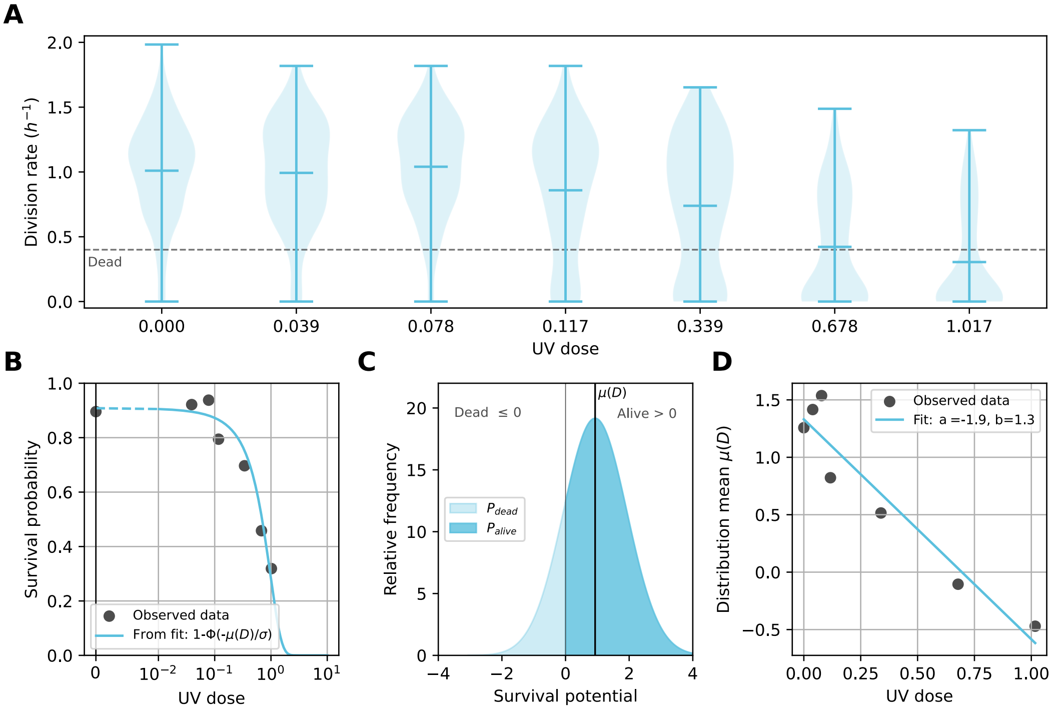


**Figure S5. Modelling the UV killing process using a statistical framework**. **A:** Classifying dead and alive cells based on growth rates recorded after UV projections (Fig. 2C). Cells with a division rate smaller than 0.4 $h^{-1}$ were considered to be dead. The UV dose $D$ is corrected for off-target effects. One unit of UV dose corresponds to $\approx$ 8.8 W $cm^{-2}$ for 300 s. **B:** Survival probabilities $P_{alive}(D)$ based on the classification from **A** (Observed data) and sigmoid fit from the cumulative distribution function of a normally distributed survival potential model. **C:** Survival potential model with dose-dependent mean $\mu(D)$. For each dose $D$, $\mu(D)$ is adjusted such that the area to the right of the origin equals the corresponding survival probability. **D:** Calculated distribution means and linear fit.

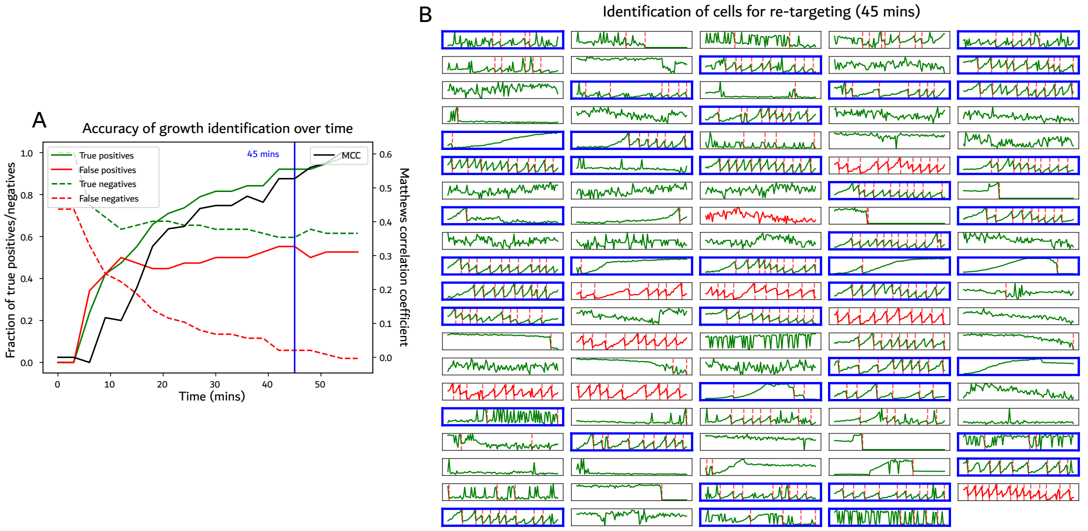


**Figure S6. Identification of cells for re-targeting**. Cells are identified as growing if the cell length $L$ is on average increasing: $L_{t+1} > L_t$ in over 50% of time points. **A:** Accuracy of identification over time. MCC = Matthews Correlation Coefficient. **B:** Cells identified for re-targeting after 45 min are shown with a blue border. Green = GFP (targeted population), red = mCherry (untargeted population).

**Table S1. Comparison of single-cell screening and selection methods applicable to bacteria.** Throughput is the approximate number of individual variants or cells that can be interrogated in a single experiment; orders of magnitude are indicative and depend strongly on implementation. “Dynamic” indicates whether each cell is measured repeatedly over time rather than at a single time point. “Iterative” indicates whether selection can be re-applied to the same physical population without re-loading or re-culturing the sample. Note that this table focuses on selection based on optical phenotypes; for review of growth-coupled phenotypes see [7].

| Method | Description | Phenotypes measured | Dynamic | Iterative | Selection method | Throughput | Refs |
|---|---|---|---|---|---|---|---|
| Well-plate screening | Variants are isolated one-per-well and assayed individually in a plate reader or colony array. | Growth, fluorescence, absorbance (bulk per well) | Yes | Yes | Physical isolation; each variant kept in a separate compartment | $10^2$–$10^3$ | [8] |
| FACS | Cells flow past a laser at high speed and are electrostatically deflected on per-cell fluorescence intensity. | Fluorescence (1–2 channels), light scatter | No | No | Physically sort desired cells into a collection vessel | $10^7$–$10^8$ | [9, 10] |
| Droplet microfluidics (FADS / AADS) | Single cells are encapsulated in pL droplets acting as microreactors, then sorted on droplet optical signal. | Fluorescence or absorbance of secreted / lysate product | Limited | No | Dielectrophoretic deflection of desired droplets | $10^7$–$10^8$ | [11–14] |
| Image-activated cell sorting (IACS) | Imaging flow cytometry with real-time image classification triggering physical sorting. | Morphology, fluorescence, sub-cellular localisation | No | No | Physically sort desired cells into a collection vessel | $\sim10^6$ | [15] |
| Digital Colony Picker (DCP) | Cells are compartmentalised in a 16,000-chamber chip, monitored by deep-learning image analysis, and exported individually by laser-induced bubble ejection. | Growth, morphology, metabolic activity | Yes | No | Contactless laser ejection of desired clones, one at a time | $\sim10^4$ | [16] |
| Optofluidic nanopen arrays (Beacon) | Single cells are positioned by light into perfused nanolitre chambers, imaged repeatedly, then exported optically. | Growth, fluorescence, secreted product titre | Yes | Yes | Optical repositioning to physically export desired clones | $10^3$–$10^4$ | [17] |
| Microcapillary arrays with laser extraction ($\mu$SCALE) | Cells are segregated in a dense glass capillary array, assayed in situ over time, and recovered by pulsed-laser ejection. | Growth, fluorescence, binding affinity, enzyme kinetics | Yes | No | Pulsed-laser extraction of desired capillary contents | $\sim10^6$ | [18] |
| Mother machine with *in situ* genotyping | Pooled libraries are imaged over many generations in trenches, then fixed and barcodes decoded by FISH or dCas9 readout. | Growth, morphology, fluorescence, sub-cellular localisation | Yes | No | None: all variants are read out and genotype–phenotype pairs recovered computationally | $10^5$–$10^6$ | [19–21] |
| Mother machine with optical tweezers | Individual cells of interest are optically trapped and physically extracted from trenches after imaging. | Growth, morphology, fluorescence | Yes | No | Physical extraction of desired cells, one at a time | $10^5$ screened, $10^1$ extracted | [22] |
| **MiPS (this work)** | **Cells are imaged and analysed in real time in the mother machine, and patterned light is projected to kill non-desired cells in place.** | **Growth, morphology, fluorescence** | **Yes** | **Yes** | **Counter-selection: photo-kill non-desired cells, recover survivors in the outflow** | $\mathbf{10^4}$**–**$\mathbf{10^5}$ | — |
